# BugBase predicts organism-level microbiome phenotypes

**DOI:** 10.1101/133462

**Authors:** Tonya Ward, Jake Larson, Jeremy Meulemans, Ben Hillmann, Joshua Lynch, Dimitri Sidiropoulos, John R. Spear, Greg Caporaso, Ran Blekhman, Rob Knight, Ryan Fink, Dan Knights

## Abstract

Shotgun metagenomics and marker gene amplicon sequencing can be used to directly measure or predict the functional repertoire of the microbiota *en masse*, but current methods do not readily estimate the functional capability of individual microorganisms. Here we present BugBase, an algorithm that predicts organism-level coverage of functional pathways as well as biologically interpretable phenotypes such as oxygen tolerance, Gram staining and pathogenic potential, within complex microbiomes using either whole-genome shotgun or marker gene sequencing data. We find BugBase’s organism-level pathway coverage predictions to be statistically higher powered than current ‘bag-of-genes’ approaches for discerning functional changes in both host-associated and environmental microbiomes.

## Background

The association of microbiota with a growing number of acute and chronic human diseases, together with decreasing costs of next-generation sequencing, has resulted in an exponential increase in the number of host-associated microbiome studies [1,2]. The microbiomes of various host body-sites have been implicated in many aspects of human health, including metabolism [3,4], immune response [5,6], disease state [7,8] and even behavior [9,10]. As the microbiome field shifts from detecting disease associations to identifying disease mechanisms, it is critical to understand the functional capacity of the microbes within a microbiome. It is also important to understand clinically relevant microbiome phenotypes when attempting to design microbiome-targeting therapies. For example, having a better understanding of the proportion of Gram negative and Gram positive microbes in the GI tract would allow for more-targeted antibiotic treatment approaches [11], and determining the anaerobe and pathogen content of donor fecal samples could help predict the efficacy and safety of fecal microbiome transplants [12,13].

Common methods to analyze bacterial microbiomes include amplicon sequencing of the 16S rRNA marker gene (16S) and whole genome sequencing (WGS) of the entire microbial metagenome. Both 16S sequencing and WGS approaches are effective for determining the taxonomic composition of a microbiome [2], with WGS also allowing better taxonomic resolution and direct profiling of functional pathways within a microbial community. Predicting the metagenome’s functional content from marker gene surveys using full reference genomes is also possible with tools such as PICRUSt (Phylogenetic Investigation of Communities by Reconstruction of Unobserved States) [14]. Regardless of sequencing approach, current methods for analyzing the functionality of a microbiome often use large microbial pathway databases such as KEGG [15] and COG [16]. Consequently, many current tools cannot use sequencing data to readily infer common microbial phenotypes, including oxygen tolerance and Gram staining, which are annotated based on experimental data regardless of pathway presence. Additionally, current outputs describing the functional capacity of a given microbiome can be overwhelming given the number of distinct genes in a metagenome, and because the biological meaning of a functional change is often lost after collapsing pathways to higher hierarchical levels. Finally, many current tools determine the functional capacity of the metagenome as a whole, implying that the genomic content of each microbe is shared across the community (Table 1). We refer to this as the bag-of-genes approach, which contrasts with inferring the functionality or phenotype of each individual strain or species, referred to here as the organism-level approach.

**Table 1.**
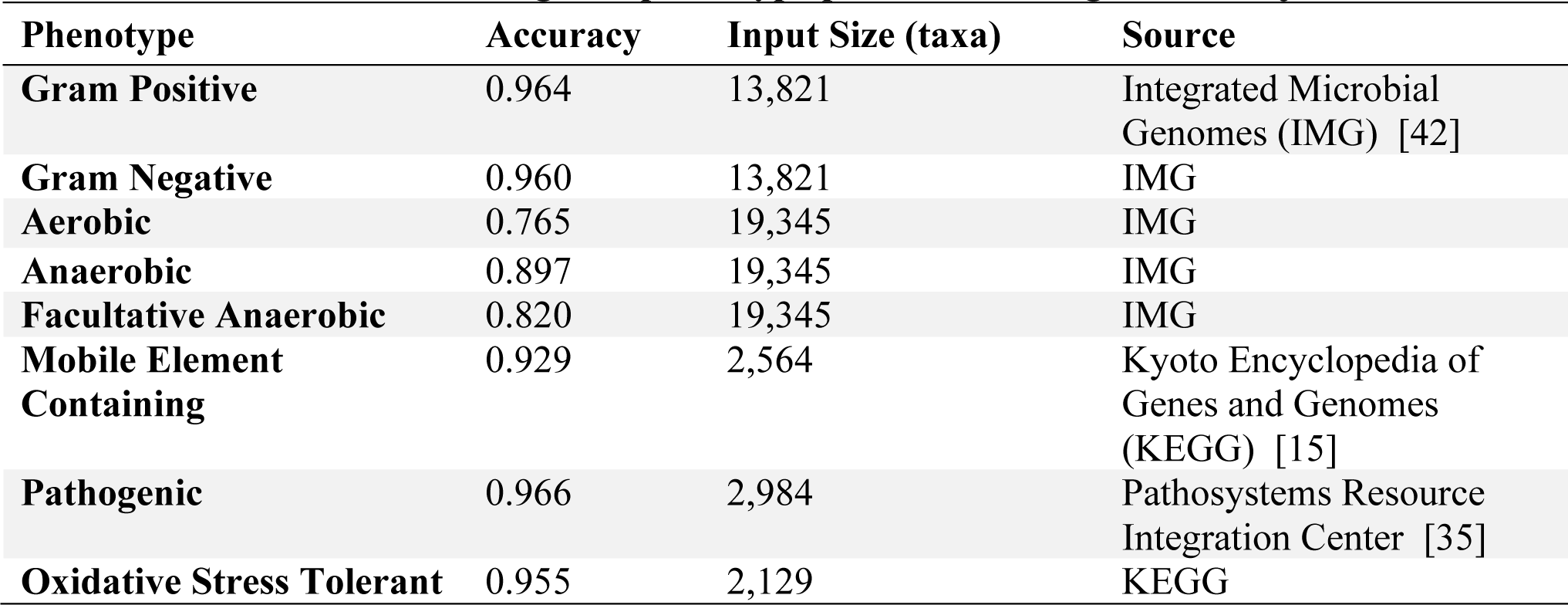
Current tools for metagenomic and trait predictions.

To solve these problems, we have developed BugBase, a novel algorithm with a user-friendly graphical user interface for analyzing microbiome data. BugBase leverages pre-existing databases, annotations and frameworks, along with manual curation, to provide users with biologically relevant microbiome phenotype predictions at the organism level. We applied BugBase to published microbiome data sets across a variety of disciplines including precision medicine, agriculture and environmental studies, demonstrating its utility for identifying interpretable biological findings not possible with current tools. Although BugBase is unable to model and predict cell-to-cell heterogeneity within strains due to genetic mutations and horizontal gene transfer, we find that organism-level predictions of functional pathway coverage are nonetheless more powerful than bag-of-genes approaches allowing detection of altered microbiome functions when bag-of-genes predictions do not. BugBase also provides predictions for simple, biologically interpretable traits such as Gram staining, oxygen tolerance, biofilm formation, pathogenicity, mobile element content and oxidative stress tolerance (Table 2), that are not available with many current tools, although traits such as mobile element content that are subject to microevolutionary processes should be interpreted with caution. BugBase works with 16S and WGS sequencing data, and can also be customized to predicted user-specified traits of interest. BugBase is available for use as a web application (http://bugbase.cs.umn.edu), and is also freely available for download (https://github.com/knights-lab/BugBase).

**Table 2.**
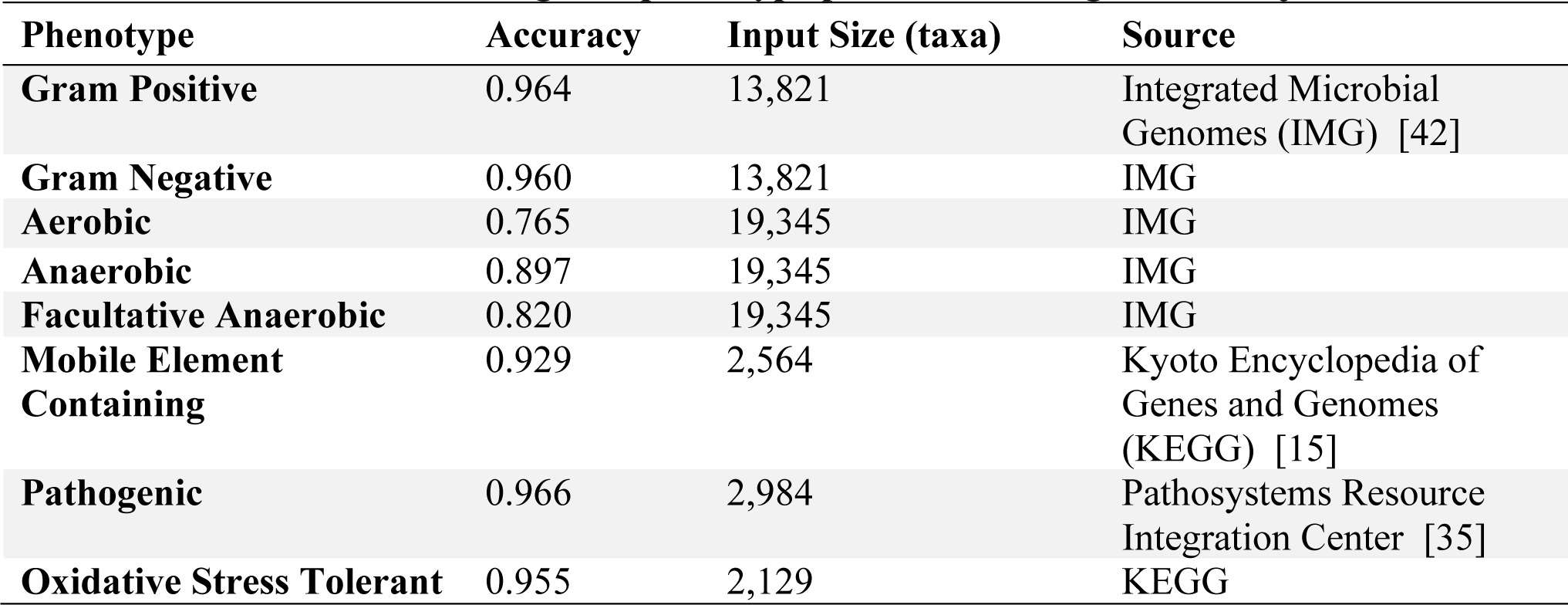
Cross-validation of BugBase phenotype predictions using a ten-fold jackknife.

## Results

### BugBase Workflow

BugBase predicts microbial phenotypes at the organism-level level (Figure 1). To do so, BugBase can use either direct observation of species and strain markers via WGS sequencing data, or a phylogenetic approach to predict genomic content of operational taxonomic units (OTUs) sequenced by 16S amplicon sequencing [14].

**Figure 1.**
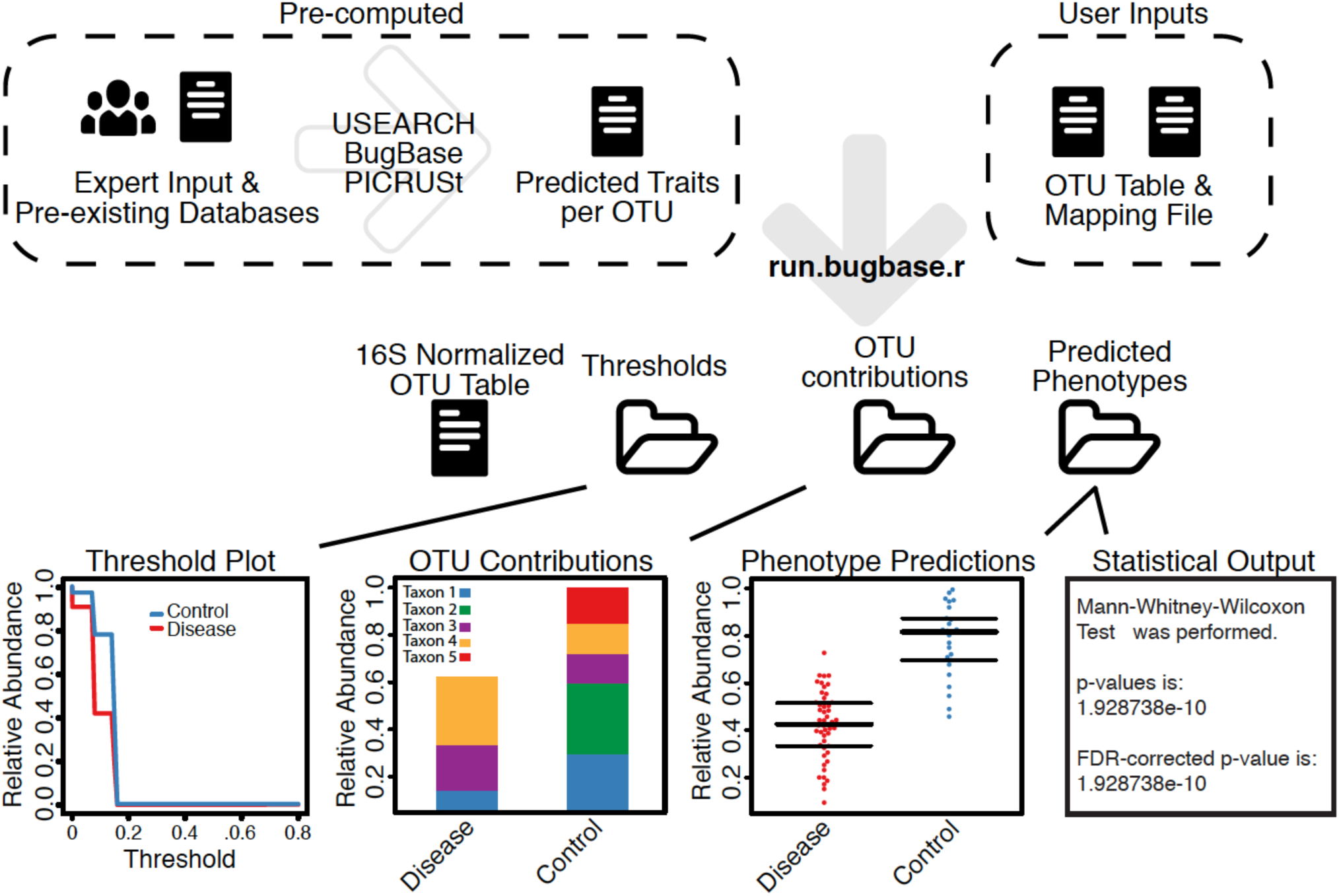
The BugBase workflow. Precomputed genomic content for reference OTUs is used to predict trait coverage for each OTU present in a biological sample. Trait abundance is calculated at different coverage thresholds across all biological samples in the data set (threshold plots). These abundance estimates are used to identify the coverage thresholds providing the highest variance across samples. OTUs in the dataset are normalized by 16S copy number and microbiome phenotypes are predicted for each trait and plotted as the trait relative abundances for each sample (phenotype relative abundances). The phenotype relative abundances are compared across experimental groups and statistical analysis outputs are provided. Discrete data types are compared using a non-parametric Mann-Whitney U or Kruskal-Wallis tests. Continuous data types are subject to a Spearman correlation test. The taxa contributing to each phenotype are also plotted as OTU contribution plots.

For WGS, species-level pathway coverage is calculated for each input sequence and each functional pathway of interest (for example, KEGG pathways), according to reference genome annotations. BugBase then automatically identifies a threshold describing the minimum fractional coverage of genes associated with a given trait that must be present within a given species for that trait to be considered present. Although in practice a user can set a predetermined coverage threshold for pathway identification, by default BugBase employs a data-driven approach to determine which coverage threshold to use for each pathway, as follows. First, trait relative abundances are estimated across the full range of coverage thresholds (0 to 1, in increments of 0.01) for every sample in a biological data set. BugBase then selects the coverage threshold with the highest variance across all samples for each trait in the user’s data. This allows for automatic determination of a pathway coverage threshold that exhibits potentially meaningful variation across samples in an unsupervised manner, without the risk of information leak from the sample metadata or experimental treatment-group labels. Once a threshold is set, BugBase then generates a final organism-level trait prediction table, which contains the predicted trait relative abundances for each sample. For both WGS and 16S data, user defined thresholds can be used to set a higher or lower stringency for phenotype prediction. For example, if the user knows that all genes in a pathway must be present for a microbe to possess a phenotype and that no other genes perform analogous functions to those in the pathway, a higher threshold (1) should be set by the user (Supplemental Figure 1). In general, a lower threshold results in higher sensitivity and lower specificity in detecting differences between groups, whereas a higher threshold results in lower sensitivity and higher specificity.

For 16S or other amplicon data, precalculated files that specify the predicted gene content or empirical trait associations for each OTU were generated using PICRUSt (Figure 1). Given a user’s OTU table, BugBase first normalizes the OTU by predicted 16S copy-number, and then predicts microbiome phenotypes using provided precalculated files and the novel approach to automatic pathway coverage estimation mentioned above. BugBase predicts that an OTU possesses a phenotype based on an empirical annotation in the cases of Gram staining and oxygen tolerance, or using an automatically identified minimum trait coverage threshold as described above for WGS data.

BugBase will optionally perform automated hypothesis testing for and visualization of differentiated traits according to user-specified metadata, and generates taxa-contribution plots depicting the relative abundances of trait-possessing taxa (Figure 1). Additional outputs include a statistical summary file that includes the results of non-parametric differentiation tests (Mann-Whitney U or Kruskal Wallis) or correlation tests (Spearman) for discrete and continuous data, respectively.

### Biological validation of predicted phenotypes in clinical and environmental microbiomes

Using five biological datasets of clinical and environmental relevance we assessed the ability of BugBase to predict interpretable organism-level phenotypes of microbiome members. These include high concordance of BugBase traits with clinical diagnostic scores in patients with bacterial vaginosis, ecologically relevant differences between major and minor human body habitats, differences in facultative anaerobes and other traits between western and non-western human gut microbiomes, and the association of oxygen tolerance with geochemical gradients in soil and water samples.

Previous studies have shown distinct differences in the distribution of taxa within the stool and oral cavity, including an overall higher proportion of Firmicutes and Bacteroidetes in stool than in the oral cavity, and a higher proportion of Actinobacteria in plaque than on the tongue dorsum [17]. Translating these differences into predicted phenotype differences is accomplished easily with BugBase. Compared to the oral cavity, BugBase predicted the stool microbiome to have a significantly higher proportion of anaerobic bacteria and a significantly lower proportion of biofilm forming bacteria (*p* <0.001). The differences in proportions of anaerobic bacteria across these body sites is well documented, and is attributed to a lack of oxygen available within the lumen of the lower gastrointestinal tract [18]. Differences in the proportion of oxygen-tolerant bacteria within the two types of plaque has also been explored, with previous reports stating an increase in anaerobic bacteria within the subgingival plaque in comparison to the supragingival plaque, due to the partially hypoxic environment below the gums [17]. BugBase readily identifies these differences, as seen by the significant difference in the proportion of anaerobic bacteria between the supra- and sub-gingival plaque (Figure 2A, *p* < 0.001). Additionally, differences in the proportion of biofilm-forming bacteria can be noted between the two plaques (*p* < 0.001), with the supragingival plaque predicted to have significantly more biofilm forming bacteria than the subgingival plaque. BugBase attributes differences in the abundance of predicted biofilm forming bacteria to the higher proportion of *Actinomyces* (phylum Actinobacteria) within the supragingival plaque samples (Figure 2B, Supplemental Figure 2), supported by previous reports [17,19].

**Figure 2.**
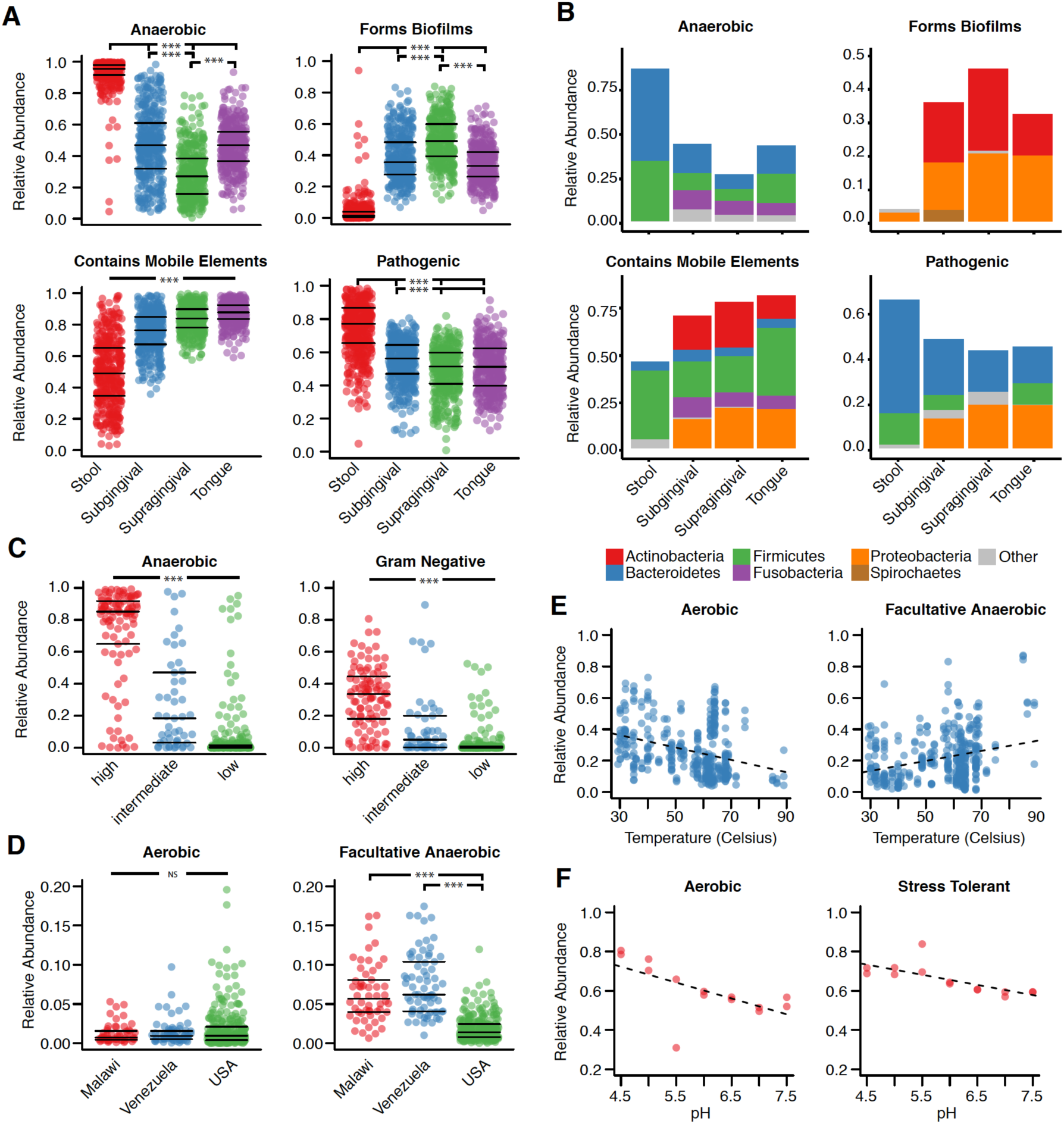
BugBase finds differences in host associated and environmental microbiomes. (A) BugBase was used to predict the proportion of anaerobic, biofilm forming, mobile element containing and pathogenic bacteria within microbiomes of stool (n=394), the tongue dorsum (n=368), sub- and supra-gingival plaque (n=373, n=376, respectively) from the Human Microbiome Project, and (B) the corresponding OTU contribution plots of the relative abundance of phyla possessing each phenotype are also shown. (C) BugBase was also used to predict biological phenotypes of vaginal microbiomes from individuals with low (n=248), intermediate (n=49) and high Nugent scores (n=97), (D) stool microbiomes from individuals in western (USA, n=263) and non-western countries (Malawi n=54, and Venezuela n=69), (E) from water microbiomes across temperature gradients in Yellowstone hot springs (n=412), and (F) soil microbiomes across pH gradients (n=14). Discrete phenotype relative abundances were compared using pair-wise Mann-Whitney U tests with false discovery rate correction, *** denotes *p* < 0.001. Continuous covariates were subject to a Spearman correlation test, *p* < 0.05 for plots shown.

BugBase also predicted the stool to have significantly more bacteria that are potentially pathogenic and significantly fewer bacteria that contain mobile elements than any of the oral cavity sites (*p* < 0.001). The high pathogenicity prediction for stool is driven mostly by the higher proportion of *Bacteroides* (phylum Bacteroidetes) in stool compared to the oral samples (Figure 2B, Supplemental Figure 2). Taxa within the *Bacteroides* genus are generally considered commensals, but like other opportunistic pathogens, such as *Enterococcus faecalis*, members of the *Bacteroides* genus often contain genes for virulence factors, including genes for enterotoxins, capsule formation and proteases [20]. Under normal homeostatic conditions *Bacteroides* species rarely express their virulence factors [20], but because BugBase’s predictions rely on genomic content, the *potential* pathogenic capabilities of the gastrointestinal microbiome are contrasted with those of the oral microbiome. The oral taxa predicted to have pathogenic potential include those previously reported as playing a role in periodontal disease, including *Porphyromonas* [21] and *Capnocytophaga* [22]. The higher relative abundance of taxa predicted to contain mobile elements in the oral samples is driven primarily by the higher relative abundance of Proteobacteria and Actinobacteria in the oral samples that house integrative and conjugative elements (a subset of mobile elements) within the Tn916 family [23]. BugBase predictions for mobile element content, however, should be interpreted with caution; recent studies have shown that although the total repertoire of mobile elements is mostly shared across geographic locations, the mobile element content can become population specific and may vary within an individual over short periods of time [24].

BugBase predictions were further validated using clinical diagnosis data for bacterial vaginosis samples [25]. Gram staining predictions of vaginal microbiome samples had high concordance with their Nugent scores, which is a standard clinical test, measuring Gram staining and bacteria by microscopy for bacterial vaginosis diagnosis (Figure 2C). BugBase predicted samples from those with high Nugent scores (bacterial vaginosis) to have significantly more anaerobes and Gram negative bacteria than those with intermediate or low Nugent scores (p > 0.001). Our analysis demonstrates that BugBase traits based on next-generation sequencing data have high concordance with a more traditional and laborious clinical test based on microscopy.

To determine the efficacy of BugBase-predicted traits for discriminating between gut microbial communities from humans living under varied lifestyles and diets, we analyzed data from a study of the fecal microbiome from individuals in villages in the Amazonas state in Venezuela, rural communities in Malawi and urban communities in the United States of America [26]. The original analysis of this dataset highlighted the drastic differences in fecal community composition between those in western and non-western countries, which was primarily driven by the predominance of *Prevotella* in the non-western samples and predominance of *Bacteroides* in the samples from the USA [26]. Using BugBase we can also see differences across the geographical locations at the phenotypic level. Unlike the differences detected across body sites in the human microbiome project (HMP) data above, no significant difference in the abundance of aerobic bacteria was seen across host country (*p* > 0.620), due to the anaerobic conditions of the lower gastrointestinal tract (Figure 2D). However, BugBase did predict those from the USA to have significantly lower levels of facultative anaerobes than those in non-western communities (Malawi and Venezuela, *p* <0.001). Also, those from the USA were predicted to have significantly fewer pathogenic bacteria and significantly more Gram positive bacteria than those in non-western communities (*p* < 0.001, Supplemental Figure 3). Although Proteobacteria is not the predominant phylum in the non-western samples, it is the taxon driving the significant differences in the microbiome phenotypes predicted (Supplemental Figure 3A). The ability to detect a non-predominant taxon driving microbiome phenotype differences highlights BugBase’s unique approach to characterizing microbiomes in a biologically relevant context. Increased levels of Proteobacteria in the stool of non-western populations has also been reported in other studies, confirming that our predictions are not an artifact of the dataset [27].

**Figure 3.**
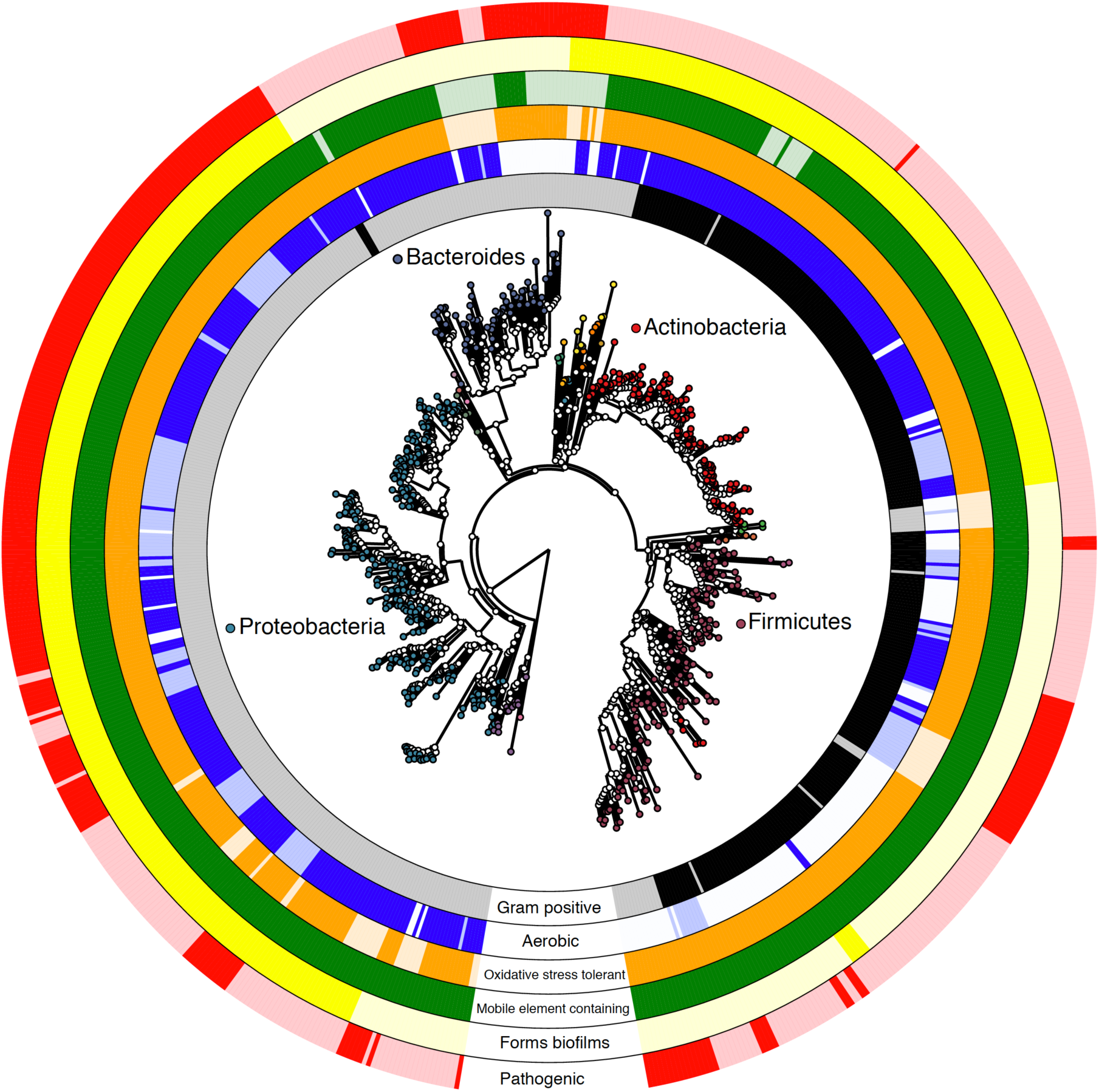
BugBase prediction across the tree of stool-associated bacteria from the Human Microbiome Project. The phylogenetic tree was created by pruning the Greengenes reference tree to tips corresponding to genera within the stool samples from the Human Microbiome Project. Tips are colored by phylum and rings are colored by the BugBase prediction for the following phenotypes: Gram positive (black), Gram negative (grey), aerobic (dark blue), anaerobic (light blue), facultative anaerobic (medium blue), oxidative stress tolerant (dark orange), mobile element containing (dark green), biofilm forming (dark yellow), and pathogenic (red). The legend for phyla not listed is shown in Supplemental Figure 7.

We also applied BugBase to a set of environmental samples characterizing hot spring microbiomes in Yellowstone National Park, Wyoming across a gradient of temperatures (Figure 2E). BugBase predicted a negative correlation with temperature and the relative abundance of aerobic bacteria (*p* < 0.001), as well as a positive correlation with temperature and the relative abundance of facultative anaerobes (*p* < 0.001). These findings are likely due to the known inverse relationship between the amount of dissolved oxygen in water and water temperature, giving facultative anaerobes a growth advantage as the amount of dissolved oxygen decreases [28–31]. The ability of BugBase to easily characterize shifts in oxygen-tolerant bacteria within microbiome samples highlights its usefulness as a resource for environmental water sampling. We also analyzed a soil microbiome dataset, which characterized the microbial community across increasing pH over a two year span [32]. The original publication and other previous studies have shown soil pH to strongly influence bacterial community diversity, and that certain taxa, such as *Acidobacteria*, are highly sensitive to shifts in pH [33]. BugBase predicted a negative correlation with increasing pH and the relative abundance of both aerobic (Figure 2F, *p* < 0.05) and oxidative stress tolerant bacteria (*p* < 0.01). Both of these results were driven by a decrease in the relative abundance of *Acidobacteria* as pH increased, confirming BugBase’s predictive power (Supplemental Figure 4).

**Figure 4.**
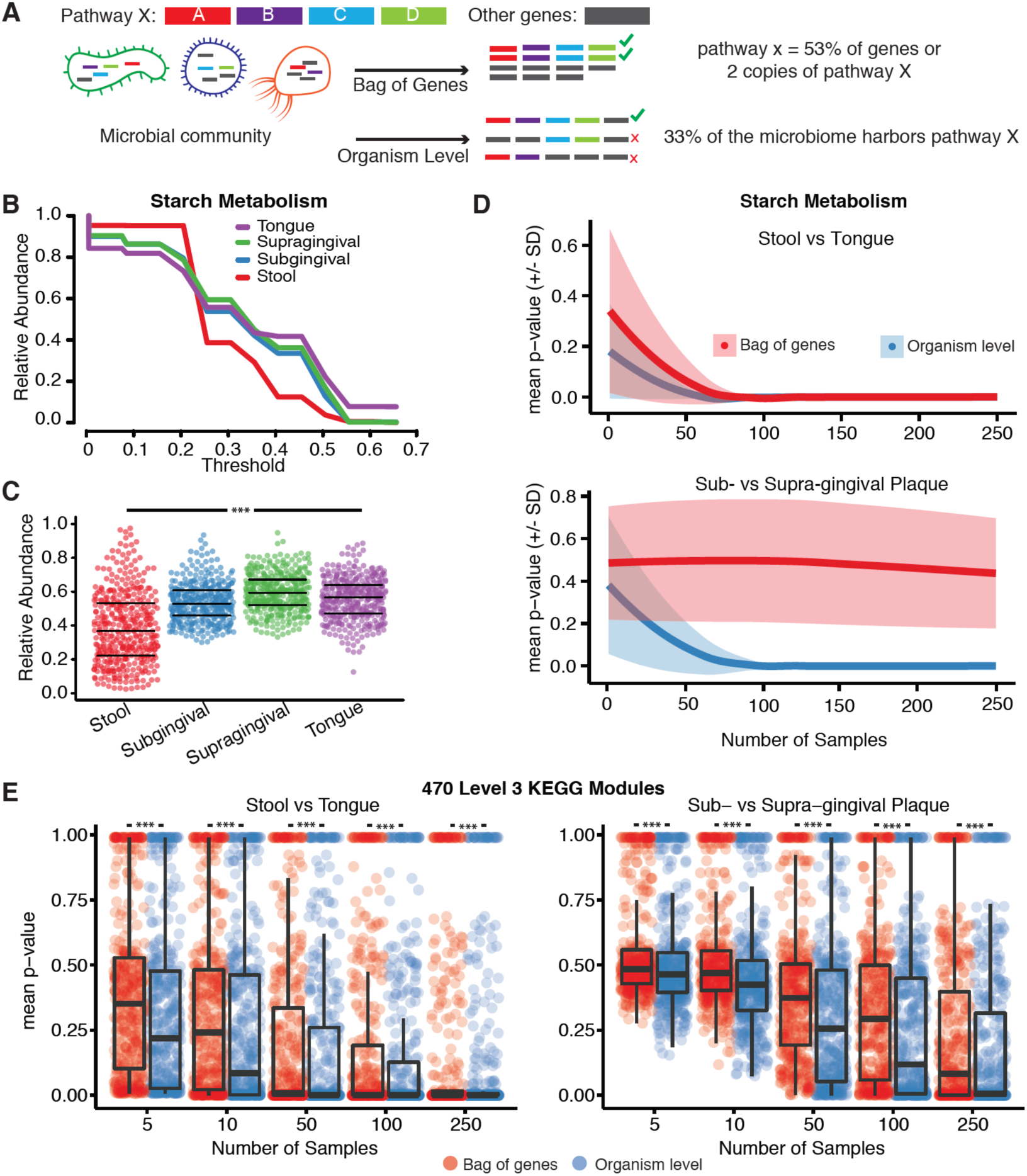
Organism-level predictions increase sensitivity. (A) BugBase’s organism-level approach versus the bag-of-genes approach used by many metagenomic analysis tools. (B) BugBase uses a threshold plot to determine the threshold (proportion of genes within the pathway) each OTU must meet to possess the starch metabolism phenotype. The threshold with the highest variance across all samples (0.25) is chosen as the default threshold by BugBase. (C) BugBase predicts significant differences in the relative abundance of starch metabolizing bacteria in the stool (n=394), the tongue (n=373) and two types of plaque (n=373, n=376). (D) The organism-level approach is more sensitive than the bag-of-genes approach at detecting changes in starch metabolism capability across taxonomically different microbiomes (stool versus tongue) at low sample numbers, and is also more sensitive to changes across taxonomically similar microbiomes (sub-versus supra-gingival plaque) at both high and low sample numbers. (E) Using 470 level-three KEGG modules, the organism-level approach is more sensitive to changes across taxonomically different (stool versus tongue) and similar microbiomes (sub-versus supra-gingival plaque). *** denotes *p* < .001 by pair-wise Mann-Whitney U tests with false discovery rate correction.

### Reference-based validation of BugBase predictions

To assess the accuracy of BugBase’s predictions we used a ten-fold cross-validation approach in which 10% of all labeled reference OTUs were withheld from the training data, and the algorithm was asked to predict traits for the held-out OTUs. This train-test process was repeated ten times ensuring OTUs were predicted by a model training only on other OTUs. BugBase’s trait predictions range in accuracy from 76% for aerobes to 97% for pathogenicity-related genome content (Table 2). Due to the limitations of ancestral state reconstruction and genome prediction, we could not assess the accuracy of BugBase’s predictions for OTUs that are not fully sequenced or have not been annotated. Because BugBase’s predictions rely on reference-genome databases for full genome prediction of OTUs, it is likely that areas of the phylogenetic tree that are sparsely populated with fully sequenced OTUs will have poorer prediction accuracy than those well populated with fully sequenced OTUs [14]. To visualize BugBase’s predictions, we have plotted BugBase’s predictions across a phylogenetic tree of taxa found in the HMP stool samples, at 97% identity of the 16S rRNA gene (Figure 3).

### Organism-level predictions are more sensitive than bag-of-genes predictions

Contrary to existing tools that use a bag-of-genes approach for metagenomic functional profiling (Table 1), BugBase predicts organism-level microbiome phenotype coverage (Figure 4A). The genome content of each OTU is evaluated to determine whether it exceeds the automatically-identified or user-set threshold for a given trait (i.e. possessing the required number of genes in the pathways involved for each trait). The relative abundance of that trait is then multiplied by the OTU counts to yield the proportion of the microbiome predicted to express the phenotype. BugBase is also unique in that users are able to take a targeted approach to functional prediction in the microbiome by specifying user-defined traits of interest. To evaluate any difference in sensitivity between the organism-level approach and bag-of-genes approach, we predicted phenotype occurrence for several well-known pathways, such as starch, glutathione, tryptophan and galactose metabolism, using the HMP dataset (Supplemental Figure 5). When comparing stool and tongue, two body sites that are significantly different in taxonomic composition, both a bag-of-genes and the organism-level approach can detect significant differences between the two body sites in a subset of functional pathways, as highlighted with the starch metabolism pathway (Figure 4B,C). As sample size decreases, however, the bag-of-genes approach becomes more affected by noise than does the organism-level approach (Figure 4D). There are also cases when only the organism-level approach reports significant differences between groups, including the comparison of supra- and sub-gingival plaque, two sample types taxonomically similar to one another (Figure 4D). In such a case, it is the within-in organism differences that cannot be detected with the bag-of-genes approach driving BugBase’s ability to predict phenotype differences between sample types. To confirm BugBase’s sensitivity, we compared 470 level-three KEGG modules across the HMP samples using a bag-of-genes approach and the organism-level approach (Figure 4E). Across a range of sample sizes, the organism-level pathway coverage predictions from BugBase were statistically higher powered than bag-of-genes approaches for discerning functional changes in host-associated and environmental microbiomes (Figure 4E).

## Discussion

BugBase users easily gain access to biological trait predictions for their WGS and marker-gene microbiome samples, accelerating the rate of scientific research through faster hypothesis generation for further experimental testing. Using a novel approach to trait-coverage estimation, the BugBase algorithm predicts organism-level microbiome phenotypes, including organism-level coverage of functional pathways and biologically-interpretable traits such as oxygen tolerance, biofilm formation capability and pathogenic capability. BugBase also allows users to predict traits not determined by most pre-existing tools, such as the proportion of the microbiome that is Gram positive and negative, as well as aerobic, anaerobic and facultative anaerobic (Table 2). BugBase’s phenotype predictions may be particularly useful for creating better-targeted clinical diagnostics and interventions. For example, bacterial vaginosis is diagnosed, in part, through microscopy of vaginal samples. This analysis requires trained microbiologists to create and score bacteria on each slide, increasing the cost and time associated with diagnosis. BugBase’s Gram staining predictions capture the differences in low, intermediate and high Nugent scores with corresponding increases in the proportion of Gram negative bacteria in the vaginal samples. Interestingly, BugBase also predicts a significant increase in the amount of anaerobic bacteria in samples with corresponding high Nugent scores (bacterial vaginosis, Figure 2C). With the rapidly decreasing costs of sequencing and quick sample-to-sequence turnarounds, BugBase’s predictions may prove to be a useful and easy diagnostic tool.

For prediction of functional pathways coverage, BugBase uses a data-driven approach to automatically detect a relevant coverage threshold for each trait based on the variability within the genomes of a given microbial community. Determining whether a given functional pathway is completely present in an organism requires precise knowledge of that organism’s genome, as well as a detailed understanding of the roles of different genes in the pathway. There can be known and unknown homologs, as well as unknown pathway components, and sharing of promiscuous enzymes across pathways. Thus, it is challenging to determine what fraction of a pathway must be identified within an OTU for the given trait to be considered present. By determining the minimum trait coverage thresholds using the observed variability in trait abundance in a given data set, rather than using a predetermined threshold, we find that we can gain important insights into traits at a coverage level that is driving the biological differences across many different types of samples.

BugBase’s predicted organism-level trait coverage also provides higher sensitivity than existing WGS or marker-gene analysis tools, and allows users to visualize changes in microbiome phenotypes across treatment groups or gradients (Figure 4). Requiring organism-level coverage of traits may be overly conservative in certain cases where bacteria are performing co-metabolism. In these cases, organism-level predictions may result in false negatives or failure to identify the potential for activity in a given pathway. However, this generally requires that the cells sharing a metabolic pathway be physically proximal in the environment, and it requires the cells to have the appropriate transporters for producing and consuming the metabolites in question. Thus, the organism-level predictions from BugBase will often be more conservative, reporting only those traits for which sufficient coverage is predicted within an individual’s genome.

Organism-level trait coverage also proves useful for phenotypes driven by rare or minor members of the microbiome, especially for WGS data. If a community contains only a small proportion of a rare but phenotypically important bacterial species, using a WGS approach in combination with a bag-of-genes analysis permits detection of only a fraction of the rare-member’s genome. This may result in being unable to detect the rare pathway of interest within the metagenome (Supplemental Figure 6). BugBase, however, requires the rare member to only be observed once for its traits to be considered present in the community. Thus, BugBase permits high resolution functional analysis of microbiomes, especially at lower WGS depths.

Due to the modularity and open-source nature of the BugBase algorithm, users can easily predict custom phenotypes by creating traits containing genes of choice at the level of sequences or homologous gene groups (Supplemental Figure 5). BugBase can also incorporate additional domains of life as reference-genome databases grow. This allows for targeted prediction of the fraction of cells containing interpretable traits in microbiome samples for hypothesis testing, although we recommend that identified traits be confirmed experimentally. BugBase leverages known reference genomes, mixed-membership strain assignment, and phylogenetic relationships to estimate genomic content of species (WGS) and OTUs (16S) for which exact reference genomes do not exist. When used with 16S data, BugBase faces the same limitations as the PICRUSt tool [14]. These include an inability to study eukaryotic and viral microbiome community members, an inability to distinguish between OTUs at the strain level in many taxa due to constraints inherent with 16S sequencing, lower prediction accuracy in branches of the phylogenetic tree with fewer annotated OTUs, and the inability to account for horizontal gene transfer between OTUs [14]. For WGS data, however, BugBase can capture strain resolution for bacteria fully sequenced, and eukaryotic and viral trait prediction can be easily added as needed. Due to the reliance of BugBase on fully sequenced organisms, it is designed to provide the basis for further hypothesis formation and exploration, and not for conclusive quantitation of organism-level traits. However, we find that BugBase provides readily interpretable biological traits and substantially improves power and interpretability in microbiome studies across a wide variety of disciplines.

## Methods

### Reference databases

BugBase uses input from various databases, such as Integrated Microbial Genomes (IMG) [34], the Kyoto Encyclopedia of Genes and Genomes (KEGG) [15] and the Pathosystems Resource Integration Center (PATRIC) [35] to categorize six main phenotype categories: Gram staining, oxygen tolerance, ability to form biofilms, mobile element content, pathogenicity and oxidative stress tolerance (Table 2, Supplemental File 1). Empirical traits predicted by BugBase are annotated based on experimental data, including Gram staining and oxygen tolerance. Empirical trait information was parsed from the IMG [34] version 4.0 database and compiled in a format needed for downstream predictions, although any combination of marker gene and reference database can be used. Non-empirical traits that are inferred based on pathway or gene presence and require expert knowledge, such as biofilm formation, oxidative stress tolerance and mobile element content were manually curated by pulling relevant open-reading frames for each trait from the KEGG database [15]. Pathogenicity data was curated using all virulence factors reported within the PATRIC databases [35]. A list of genes for each non-empirical traits can be found in Supplemental File 1. Alignments of each open-reading frame against all genomes within IMG were performed with USEARCH, using the ‘usearch_global’ parameter with an ID match of 0.6 and a query coverage of 0.8 [36]. Outputs were parsed and formatted as tab-delimited text files denoting the gene count for each strain within IMG. For 16S data analysis, the resulting files were used in PICRUSt’s genome prediction pipeline, which predicts the occurrence of a given gene or empirical trait for all Greengenes OTUs [37], including those with incomplete genome sequencing, based on phylogeny and the genomic content of fully sequenced IMG reference genomes [14]. For these predictions we used the Greengenes 97% OTU tree and ancestral state prediction feature of PICRUSt. The predicted precalculated files are calculated offline to avoid unnecessary calculations during microbiome phenotype predictions. For WGS, precalculated files were generated using all organism-specific data from KEGG [15] and IMG [34] for pathway and empirical traits, respectively.

### Predicting organism-level phenotypes

To predict microbiome phenotypes, BugBase takes as input either a biom-formatted OTU table (JSON version) [38], for 16S data, or a QIIME-formatted OTU table (biom or classic) for WGS data. Optionally, the user may supply a QIIME-compatible mapping file containing metadata for each sample. For 16S data, the OTU table is normalized by 16S copy-number prior to trait prediction. The taxon table (WGS strain table or normalized 16S OTU table) is defined as *A* = *a*_*i j*_ ∈ ℝ ^*m×n*^, with relative abundance *a*_*i j*_ of taxon *j* in sample *i*. The precalculated trait coverage table is defined as *P* = *p*_*j k*_ ∈ ℝ^*n × p*^, with relative abundance *p*_*j k*_ of trait *k* in taxon *j*. For a particular trait *k*, we then estimate the fraction of organisms *q* in each sample *i* with at least fractional trait coverage *t*_*l*_. If a user-defined threshold is not set, we sweep across a range of trait coverage thresholds *t* = {0.01,0.02, …,1} to obtain a 3-dimensional array *Q* = (*q*_*i k l*_) ∈ ℝ^*m*×*p*×100^. For each trait and threshold, we calculate the variance of that trait’s relative abundance across all samples in the data sets var(*Q*_·*k l*_), and then choose the threshold 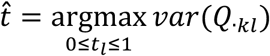 that maximizes this variance. The dependence of trait relative abundances on minimum trait coverage thresholds may be visualized using trait coverage plots as shown in Figure 1. Trait coverage plots and their corresponding tabular outputs depict the predicted fraction of cells in the microbiome (*y* axis) with a given fraction of trait coverage (*x* axis). After selecting a coverage threshold we then collapse the 3-dimensional array to the final 2-dimensional sample-trait table *T* = (*t*_*i k*_) ∈ ℝ^*m× p*^. Users also have the option to use the coefficient of variation, instead of variance, as the threshold-determining metric.

### Assessment of accuracy

The phenotype prediction accuracy of BugBase for 16S-based predictions was measured with a 10-fold cross validation strategy using empirical trait occurrence and known gene content for non-empirical traits from the original USEARCH alignments. WGS-based predictions are expected to be uniformly higher in accuracy due to the direct mapping of sequences to strains. Test and training files were created that consisted of 10% of the labeled reference OTUs (fully sequenced) and all other sequenced OTUs, respectively. Predicted gene content or trait annotation was compared to the known gene content or trait annotation for each OTU. The train-test process was repeated ten times to ensure OTUs were predicted by a model training only on other OTUs. The proportions of accurate predictions are reported in Table 2. For non-empirical traits, the accuracy metric reported is for the prediction of gene occurrence in the predicted versus known sample. The presence of non-empirical phenotypes for each OTU would need to be tested experimentally in order to assess true phenotype prediction accuracy.

### Greengenes 97% OTU Phenotype Prediction

Phenotype predictions for stool OTUs in the Greengenes 97% OTU dataset [37] were parsed from BugBase’s precalculated files. We used the threshold values generated by BugBase’s default setting from comparing the stool, tongue, sub- and supra-gingival plaque from the Human Microbiome Project analysis mentioned below. The phylogenetic tree was generated with GraPhlAn v0.9 [39].

### Phenotype Predictions for Published Datasets

Sequences were downloaded from the Human Microbiome Project website (http://hmpdacc.org/resources/data_browser.php) and OTUs were assigned using the default settings in QIIME v1.8.0 [40], using a closed reference approach at 97% identity against the 97% OTU Greengenes representative set (v13.5.8) [37]. The resulting OTU table and mapping file were parsed to include only samples with at least 500 sequence counts from tongue dorsum (n=368), stool (n=394), sub-(n=373) and supra-gingival plaque (n=376), using QIIME v1.8.0. Data for the western versus non-western comparisons, Yellowstone National Park hot springs predictions, and soil phenotype predictions were downloaded from the Qiita website (http://qiita.microbio.me) and analyzed using QIIME v1.8.0 with the same minimum sequence depth per sample mentioned above. For the western versus non-western comparison, the OTU table and mapping file were parsed to include only samples from non-infants (>3 years old) residing in the USA (n=263), Malawi (n=54) or Venezuela (n=69). For the Yellowstone data, the resulting OTU tables and mapping files were parsed to include only water samples (n=412). Sequences from the vaginal microbiome study [25] were downloaded from the National Center for Biotechnology Information Short Read Archive (SRA022855), and OTUs were picked using NINJA-OPS [41] and a closed reference approach at 97% identity against the 97% OTU Greengenes representative set (v13.5.8) [37]. Samples with at least 500 sequence counts were retained for the final anlysis (low Nugent score, n =248; intermediate Nugent score, n=49; high Nugent score, n= 97). BugBase analysis of the filtered OTU tables and mapping files was performed using the run.bugbase.r command.

### Organism-level and bag-of-genes prediction comparisons

KEGG IDs [15] for genes involved in each pathway were used to parse PICRUSt’s precalculated file (KEGG ID counts per Greengenes OTU). These trait files were then used to predict pathway occurrence for subsets of the HMP data (stool, tongue, supra- and sub-gingival plaque), ranging from 1 sample per body site to 250 samples per body site. This was repeated 30 times per sample size, with a Mann-Whitney U test for differences in pathway abundance between two body sites completed for each repeat. The average *p*-value (loess smoothed) with standard deviations was plotted for each sample size (Figure 4, Supplemental Figure 5). A similar approach was used to compare the bag-of-genes approach to the organism-level approach for all level three KEGG modules. The mean *p*-values from Mann-Whitney U comparisons of KEGG module relative abundance differences between body sites (with 30 subsampling repeats) were plotted for the 470 level-three KEGG modules found in the tongue, stool, supra- and sub-gingival plaque from the HMP dataset (Figure 4).

## Declarations

### Availability of Data materials

BugBase can be used as a web application (http://bugbase.cs.umn.edu) and is also available as an open-source package (http://knightslab.org/tools). When used to predict the default traits or KEGG level-three modules that come with BugBase, the only dependencies required for BugBase include R and required R libraries. The full analysis workflow for the data presented here, including the data files required, is found at http://github.com/TonyaWard/BugBase_Manuscript.

### Author contributions

D.K. and T.W. conceived the method. T.W., D.K., J.M., B.H. and J.L. implemented the software. J.L., T.W. and B.H implemented the web application. D.S, R.F. and T.W. curated default pathway information and annotations. J.S., G.C., and R.K. contributed experimental data. T.W. analyzed the data. T.W., and D.K. wrote the manuscript with help from J.L., J.M, J.L., D.S., J.S. G.C., R.B., R.K., and R.F.

## Acknowledgements

We would like to thank the authors of the original studies that provided data for our analysis, and the Yellowstone Center for Resources for a research permit granted to J.R.S. to work at selected locations within the Park (permit #05664).

